# Age-associated reduction of nuclear shape dynamics in excitatory neurons of the visual cortex

**DOI:** 10.1101/2022.08.22.504704

**Authors:** Tanita Frey, Tomonari Murakami, Koichiro Maki, Takumi Kawaue, Ayaka Sugai, Naotaka Nakazawa, Taiji Adachi, Mineko Kengaku, Kenichi Ohki, Yukiko Gotoh, Yusuke Kishi

**Author notes:** Correspondence (Y.K.).

## Abstract

Neurons decline in their functionality over time, and age-related neuronal alterations are associated with phenotypes of neurodegenerative diseases. In non-neural tissues, an infolded nuclear shape has been proposed as a hallmark of aged cells and neurons with infolded nuclei have also been reported to be associated with neuronal activity. Here, we perform time-lapse imaging in the visual cortex of *Nex-Cre;SUN1-GFP* mice. Nuclear infolding was observed within 15 minutes of stimulation in young nuclei, while the aged nuclei were already infolded pre-stimulation and showed reduced dynamics of the morphology. In young nuclei, the depletion of the stimuli restored the nucleus to a spherical shape and reduced the dynamic behavior, suggesting that nuclear infolding is a reversible process. We also found the aged nucleus to be stiffer than the young one, further relating to the age-associated loss of nuclear shape dynamics. We reveal temporal changes in the nuclear shape upon external stimulation and observe that these morphological dynamics decrease with age.

## Introduction

Aging is associated with a decline in brain function, such as memory and cognition, and several neurodegenerative disorders, such as Alzheimer’s disease, Huntington’s disease (HD), and Parkinson’s disease (PD) ^1–3^. Identifying the mechanisms of brain aging is necessary for treating and preventing aging-related conditions.

Among several hallmarks of the aging process, abnormalities of nuclear properties are common features of naturally aged and senescent cells in non-neural tissues ^4–6^. In neural tissues, the abnormal nuclear shape is observed in aging-associated neurodegenerative diseases. In human patients with PD, neuronal nuclei are rather infolded, marked by their invaginated nuclear membrane, in contrast to the spherical shape observed in the hippocampus, frontal cortex, and substantia nigra ^7,8^. Furthermore, in PD-model mice, with knockout or G2019S point mutation of the Lrrk2 kinase, the nuclei in the substantia nigra and dorsal thalamus showed more infolded nuclei ^7,9^. Similar infolded nuclear shapes in neurons were found in human patients with HD and the mouse model, overexpressing *Htt* genes with expanded CAG trinucleotide ^10,11^. These infolded nuclei in neurodegenerative diseases, such as PD and HD, showed defects in nuclear compartmentalization, and a more permeable nuclear membrane ^7,10^. Nucleocytoplasmic transport of RNA was also impaired in neurons of the model mouse of HD ^11^. These reports indicate the role of nuclear properties, such as nuclear shape, in the decline of neuronal function in neurodegenerative diseases; however, it is still unknown how the nuclear properties of neurons change over the time-dependent natural aging process.

Nuclei in neurons also change their shape upon neuronal activity. Internal and external stimuli mediate neuronal activity and functional neuronal networks by inducing morphological changes in axons, dendrites, and synapses ^12,13^ and also by remodeling transcriptome during neuronal activity ^14^. In the neuronal culture of the hippocampus and dorsal thalamus, neuronal activity induced via blocking GABA receptors using bicuculline increased the portion of infolded nuclei ^9,15,16^. Also, the physiological stimulation by exposure to an enriched environment increased the amount of infolded nuclei in the CA1 of the hippocampus ^16^. Nuclear infolding in neurons induced by neuronal activity is implicated in regulating gene transcription and nuclear compartmentalization. The neurons with infolded nuclei tended to have more phosphorylated histone H3 at serine 10, a histone modification for gene activation ^15^. Satb2, a neuronal transcription factor, is also involved in inducing the infolding of neuronal nuclei ^16^, and it has been reported that multiple nuclear compartments separated by the invagination of nuclear membranes present different amounts of calcium signaling ^15^. An examination of the dynamics of the nuclear shape upon external stimuli in the physiological condition would further clarify our understanding of nuclear infolding and its role as would the age-associated differences in the response of the nuclear shape upon external stimulation, which is currently completely unknown.

In this study, we analyzed the nuclear shape of excitatory neurons in the visual cortex of young and aged mice upon light stimulation after dark rearing. Previous reports have shown that excitatory neurons show nuclear infolding upon external stimuli ^15,16^. We first confirmed the increased number of infolded nuclei in the stimulated visual cortex of young mice. To further reveal the dynamics of the infolding process in young mice, we performed time-lapse imaging of the nuclear shape of excitatory neurons in the upper layer of the visual cortex of *Nex-Cre;SUN1-GFP* transgenic mice, whose nuclei in excitatory neurons were labeled with a GFP fused with SUN1 outer nuclear membrane protein ^17,18^. Over the course of light stimulation, nuclear shapes in the excitatory neurons of the visual cortex showed dynamic changes toward an infolded morphology, which was observed 15 minutes after visual cue exposure. After turning off the light stimulation, the nuclei’s dynamics turned down, and the spherical shape reappeared, and was similar to conditions before the light stimulation, suggesting reversible changes in the nuclear morphology. Additionally, we also examined nuclear infolding in the aged brain and found that the aged visual cortex (over 2 years), even without visual stimulation, showed a higher portion of infolded nuclei than the young one. Time-lapse imaging of aged mice revealed that the nuclei showed less frequent nuclear infolding upon visual stimulation, and atomic force microscopy (AFM) analysis revealed aged neurons had stiffer nuclei. In our research, we first presented the temporal output of visual stimulation on the nuclear shape in the visual cortex of excitatory neurons and then the decline in the nuclear shape dynamics during the natural aging process.

## Results

### Infolding of nuclear shape within excitatory neurons increases upon visual stimulation in the visual cortex

Previous studies have examined the changes in nuclear shape via different evoked stimulation systems ^9,15,16^. To study the in vivo nuclear shape behavior in relation to neuronal activity upon physiological stimulation, we utilized the visual system because of its simplified manipulative approach via ocular enucleation and dark rearing. We performed enucleation of the left eye, and the mice were exposed to five days of dark rearing to reduce the neuronal activity in the visual cortex to a baseline ^19^. After 4 hours of light exposure, we confirmed the upregulation of *Npas4,* which is a representative immediate early genes (IEGs) induced by neuronal activity by reverse–transcribed quantitative PCR (RT-qPCR) analysis (Supplementary Fig. 1A, B).

Using the visual manipulation, we aimed to examine the nuclear shape in excitatory neurons of the visual cortex in *NexCre;SUN1-GFP* mice, which express the GFP-fused SUN1 protein, a protein on the outer nuclear membrane, under the control of the *NeuroD6/Nex* gene locus ^17,18^. We used 6-to 9-week-old mice, in which the critical period of visual cortex development had finished, to rule out any effects due to the particular behavior of neurons during this period ^20,21^. The left eyes of the 6-to 9-week-old *NexCre;SUN1-GFP* mice were enucleated, and the mice were kept in a dark room for 5 days and afterward were exposed to the light for 4 hours (Fig. 1A). The brain slices were immunostained with the antibodies for GFP and c-Fos to evaluate neuronal activity (Fig. 1B). The ipsilateral hemispheres of the removed left eye, which received light stimulation from the intact right eye, showed an increase in their c-Fos expression in the visual cortex in comparison to the contralateral hemisphere, which did not receive light stimulation (Fig. 1B, D). This supports the notion that our dark rearing of monocular mice and light stimulation appropriately manipulate neuronal activity. The GFP staining pattern showed spherical (arrowheads) or infolded (arrows) nuclear shapes (Fig. 1B, C), and their numbers were counted within the stimulated and deprived hemisphere of monocular enucleated mice (Fig. 1D). In the contralateral hemisphere, 60% of total nuclei showed an infolded shape. By contrast, in the ipsilateral hemisphere, the nuclear shape distribution showed a strong dominance of infolded nuclei (79%) in comparison to spherical nuclei (20%). Moreover, most of the infolded nuclei (74%) in the ipsilateral hemisphere were c-Fos-positive, suggesting that physiological visual stimulation induces nuclear infolding in the visual cortex.

**Figure 1.**
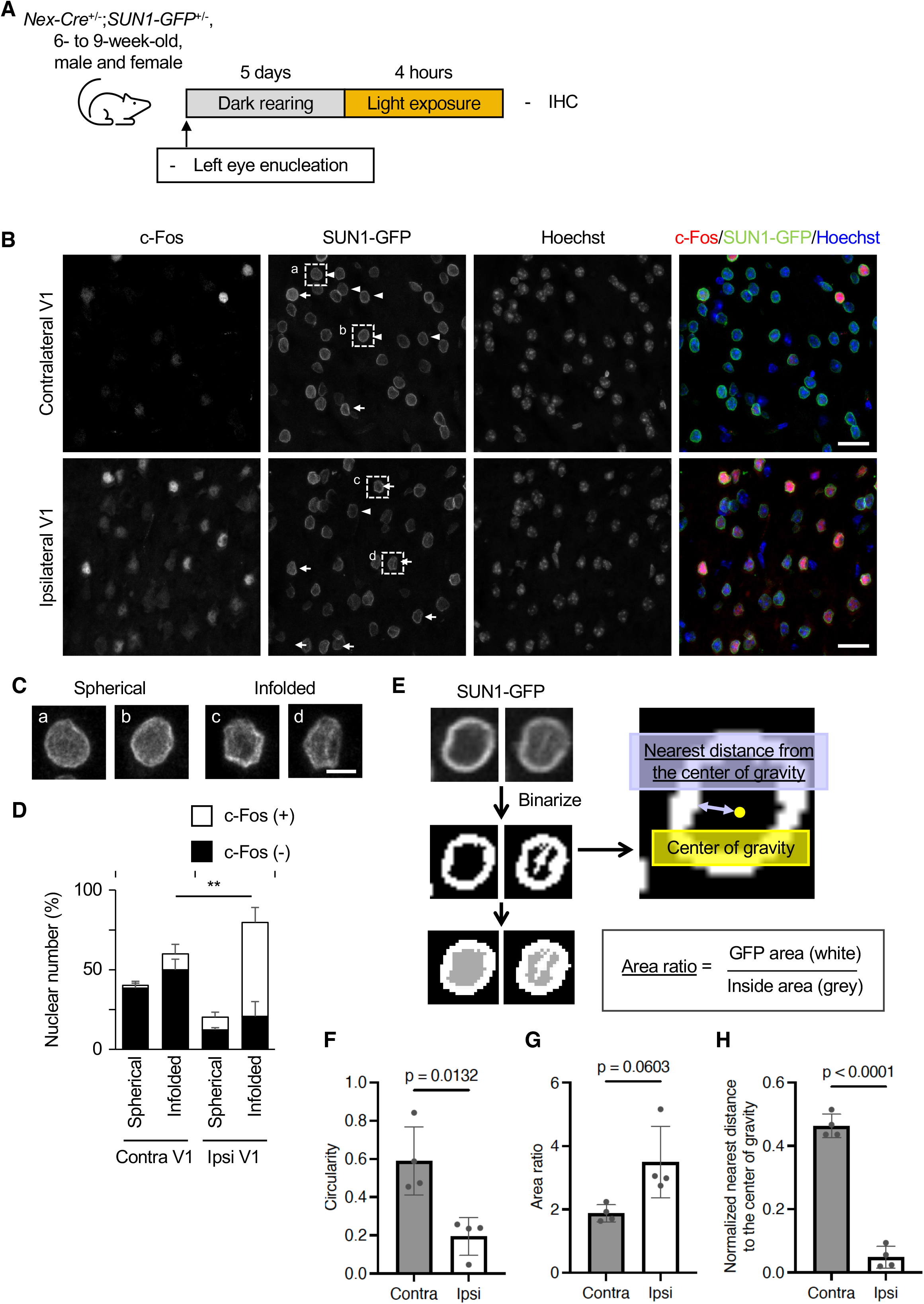
Nuclear infolding in the excitatory neurons in the primary visual cortex induced by visual stimulation. **(A)** Left eyes of 6- to 9-week-old *Nex-Cre;SUN1-GFP* mice were enucleated, and the mice were kept in a dark room for 5 days. They were stimulated by ambient light for 4 hours and subjected to immunohistochemistry. **(B)** Coronal sections of the brain were stained with antibodies to c-Fos and GFP. Nuclei were counterstained with Hoechst 33342. The images were obtained from layer 2/3 of the primary visual cortex. Arrows and arrowheads indicate infolded and spherical nuclei, respectively. Scale bars, 20 μm. **(C)** Higher magnification images of spherical and infolded nuclei indicated in (B). Scale bars, 5 μm. **(D)** Quantification of the proportion of c-Fos–positive and spherical or infolded nuclear shapes among all GFP-positive cells in all layers of the primary visual cortex. 1,842 (contralateral) and 2,926 (ipsilateral) nuclei from four independent experiments were analyzed. **(E)** Area ratio values were determined as the area of GFP signals (white) divided by the inside area (gray) after binarization of the nuclear image. The nearest distance from the center of gravity was determined as the distance from the center of gravity to the closest GFP signal. (**F–H**) Quantification of circularity (F) or area ratio (G) value of SUN1-GFP signals or nearest distance from the center of gravity (H) in the ipsilateral and contralateral visual cortex. 306 (contralateral, circularity), 291 (ipsilateral, circularity), 165 (contralateral, area ratio), 190 (ipsilateral, area ratio), 158 (contralateral, nearest distance), and 194 (ipsilateral, nearest distance) nuclei from four independent experiments were analyzed. Data are means ± s.d. *p* values were determined by two-tailed Welch’s t-test.

Since distinguishing nuclear shapes relied on the subjective observations of the SUN1-GFP signal, we next evaluated the nuclear shape using three objective values: circularity, area ratio, and nearest distance from the center of gravity. Previous studies have evaluated nuclear infolding by reduction of circularity value ^9,10^. Therefore, we first examined the circularity value of the inner area of the SUN1-GFP signal (a gray area in Fig. 1E) and found that circularity in the ipsilateral visual cortex was lower than that in the contralateral one (Fig. 1F).

Along with the outer shape changes, we noticed that some nuclei presented the SUN1-GFP signal inside the nuclei. Therefore, we also defined an area ratio value: the SUN1-GFP signal area (a white area in Fig. 1E) divided by the non–SUN1-GFP occupied area (a gray area in Fig. 1E) within the nucleus (Fig. 1E). This means that a higher area ratio value represents a more infolded or non-spherical nuclear shape. When we examined the area ratio value in the ipsilateral visual cortex, it tended to be higher than that in the contralateral one (Fig. 1G).

The changes in circularity and area ratio did not necessarily reflect the invagination of the nucleus. Therefore, we further analyzed the distance between the center of gravity (a yellow point in Fig. 1E) to the closest signal of GFP (a blue arrow in Fig. 1E). The distance to the center of gravity in the stimulated ipsilateral cortex was shorter than that in the contralateral cortex (Fig. 1H). Taken together, these results suggest that the neuronal activity by light stimulation induced the shift of nuclear shapes towards an infolded shape within excitatory neurons of the visual cortex, and that this is a novel context of nuclear infolding of neurons induced by physiological stimulation.

### The nuclear shape of excitatory neurons dynamically changes in response to visual stimulation in vivo

Since nuclear infolding can be observed upon exposure to visual stimulation in excitatory neurons within the visual cortex, this raises the question of the infolding process and its temporal relationship to neuronal activity. Therefore, we performed live imaging of the upper layer of the visual cortex in *NexCre;SUN1-GFP* mice in vivo with two-photon microscopy during visual light exposure following the prior preparations of craniotomy on the visual cortex for time-lapse imaging (Fig. 2A). After the craniotomy, the mice recovered for more than 2 weeks and then underwent harvesting in a dark room for 5 days. Then, for the “before light stimulation” recordings, using low-dose anesthesia, we placed the mice in front of a black monitor displaying a black image for approximately 1 hour. We considered this recording as a direct control for the following stimulation conditions in the same individual. For visual stimulation, the mice were exposed to spatially randomized visual cues presented on the monitor, separated by 4 seconds of gray screen presentations. Imaging of SUN1-GFP signals was displayed for approximately 20 minutes before the light stimulation showed a stable and spherical nuclear shape of excitatory neurons in the upper layer, indicated by arrows in Fig. 2B and Supplementary Video 1. However, the nuclear shape changed dynamically and became infolded during the light stimulation (Fig. 2B, Supplementary Video 2). We classified the nuclear shapes into changed or maintained and spherical or infolded before and during the exposure to light stimulation (Fig. 2C). We found that the majority of the nuclei (50%) maintained their spherical shape before stimulation, while 37% of the nuclei during stimulation changed their nuclear shape from spherical to infolded.

**Figure 2.**
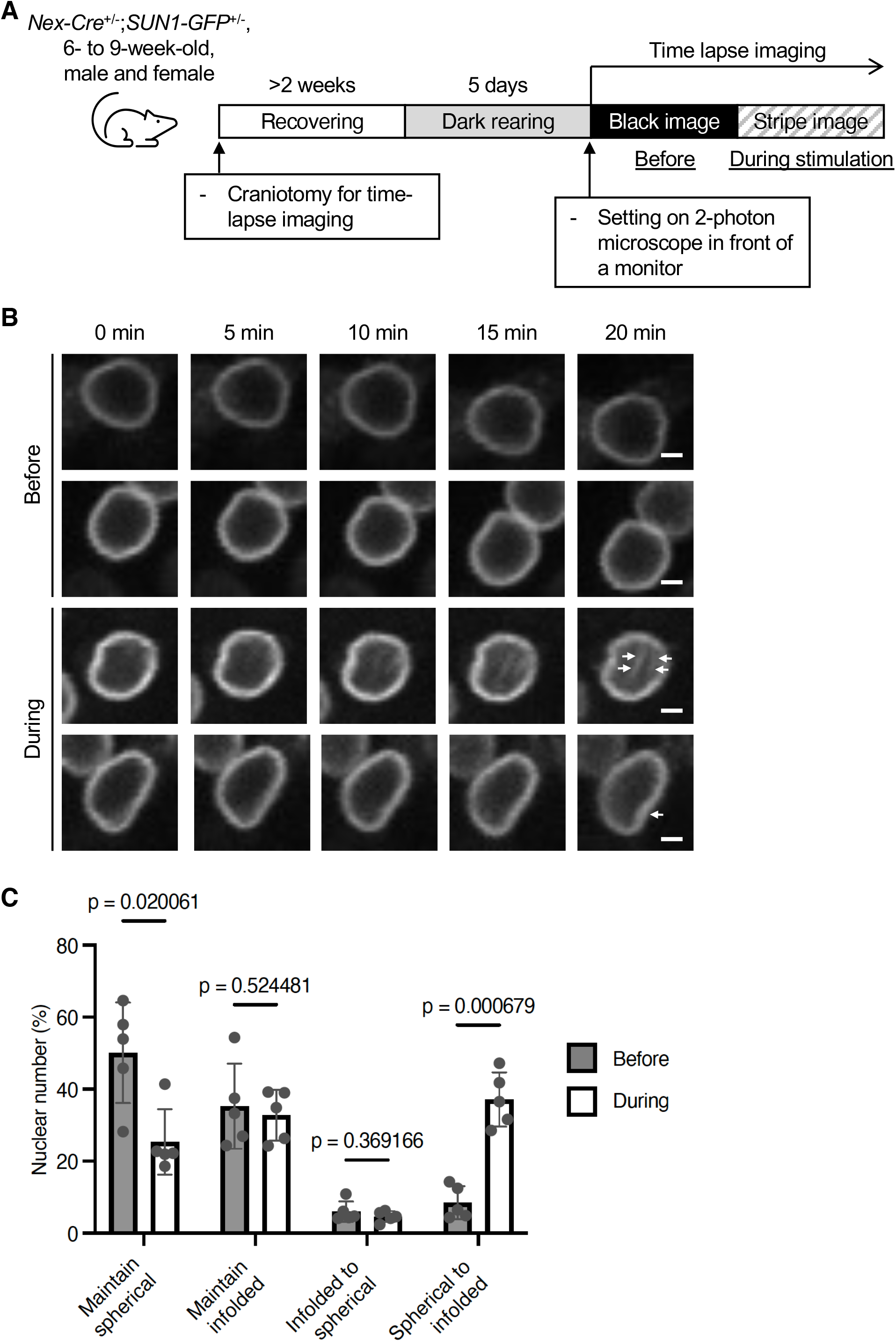
Time-lapse imaging of the nuclear shape of the excitatory neurons in the primary visual cortex upon visual stimulation. **(A)** 6- to 9-week-old *Nex-Cre;SUN1-GFP* mice were subjected to craniotomy for time-lapse imaging. After recovering for approximately 2 weeks and being kept in the darkroom for 5 days, the mice were set on the two-photon microscopy in front of the monitor. The nuclear shape visualized by GFP in layer 2/3 was recorded at an interval of 30 seconds with the black image (“before”) and then with the striped image (“during”). **(B)** Representative images of GFP signals in “before” (upper two panels) and “during” (lower two panels) conditions. Arrows indicate and invaginated area at 20 minutes. Scale bars, 2 μm. **(C)** Quantification of the proportion of the nuclei showing the “maintained” or “changed” shape between spherical and infolded during the recording are shown. 322 nuclei in “before” and 305 nuclei in “during” from five independent experiments were analyzed. Data are means ± s.d. *p* values were determined by two-tailed Welch’s t-test.

For objective evaluation of nuclear shape, we analyzed the area ratio value defined in Fig. 1E. Before light stimulation, the area ratio did not change for 20 minutes of non-light exposure (Fig. 3A). However, during light stimulation, the area ratio started to increase after 15 minutes of exposure (Fig. 3B). This suggests that the nuclear shapes of excitatory neurons in the visual cortex require approximately 15 minutes to start altering their shape upon visual stimulation. As immediate responses after stimulation for neurons, MAP kinases are activated within 5 minutes, and they expressed IEGs within 20 minutes ^22^. Our results indicate that nuclear infolding is another immediate response of neurons upon external stimulation.

**Figure 3.**
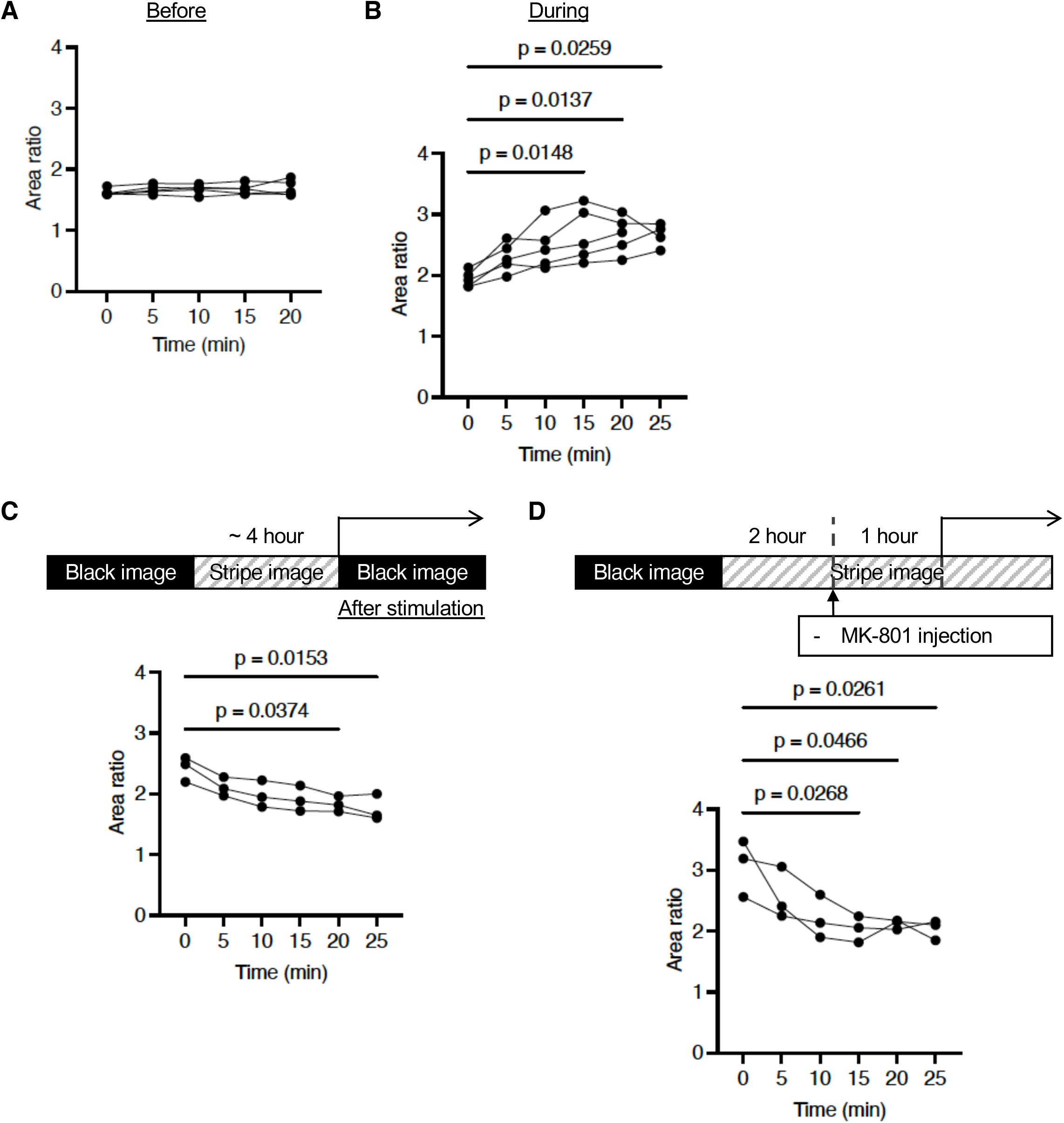
The behavior of the nuclear shape before, during, and after visual stimulation. **(A, B)** Quantification of area ratio values at indicated time in “before” (A) or “during” (B) recording. **(C, D)** Experimental schemes and quantification of area ratio values at indicated time in “after” (C) or “MK-801” (D) recording. 52 nuclei from five independent experiments (A), 38 nuclei from five independent experiments (B), 25 nuclei from three independent experiments (C), and 30 nuclei from three independent experiments (D) were analyzed. *p* values were determined by one-way ANOVA followed by Tukey’s multiple-comparison test.

### Dynamic behavior in nuclear shapes decreases after neuronal activity in excitatory neurons is inhibited

The strong relationship between neuronal activity and nuclear shape raises the question of what would happen to the nuclear shape once the neuronal activity is inhibited. First, we examined the changes in SUN1-GFP patterns after turning off the visual stimulation by switching to the black image on the monitor. During the first 20 minutes, the dynamic behavior of nuclear shapes gradually decreased and at 25 minutes became similar to the pre-stimulation area ratio value (Fig. 3C, Supplementary Video 3), suggesting that the nuclear shape takes approximately 20 minutes to inhibit the dynamics observed during stimulation.

Additionally, we also examined another method by which to inhibit the effect of neuronal activity—pharmacological inhibition of calcium influx. MK-801 is an NMDA receptor antagonist, and its administration reduces calcium influx and neuronal activity. To confirm that MK-801 interferes with neuronal activity, the expression patterns of IEG in 6-week-old *C57BL6/J* wild-type mice were analyzed. Following the 5 days of dark-rearing treatment, the mice were exposed to light for 2 hours and then injected with MK-801. RT-qPCR analysis of the visual cortex showed the weakened expression of *c-Fos* in brains injected with MK-801 compared to those injected with saline, confirming the effect of MK-801 on the neuronal activity induced by the visual stimulation (Supplementary Fig. 2A). On this basis, we performed time-lapse imaging to evaluate the effect of MK-801 on the visual cortex of *NexCre;SUN1-GFP* mice following 2 hours of light exposure and examined the response of the nuclear shape upon the NMDA inhibitor application while presenting visual cues. We imaged the visual cortex approximately 1 hour after injection and found that the nuclear shape had changed and the area ratio decreased despite the presence of light stimulation (Fig. 3D, Supplementary Video 4). These results suggest that NMDA activation and the calcium influx in excitatory neurons of the visual cortex are necessary for the dynamic changes in nuclear shape.

Taken together, these results indicate that the dynamic behavior of nuclear shapes induced by neuronal activity is a reversible event in an in vivo physiological system.

### The nuclei in aged mice are more infolded compared to those in young mice

In addition to the neuronal activity–induced nuclear infolding, previous studies have shown more infolded nuclei in human patients with PD and HD and PD and HD model mice ^7–11^. Since both PD and HD are neurodegenerative diseases associated with aging, we next examined the shape and dynamics of the nuclei in aged mice. We prepared 7-week-old and 133-week-old *C57BL6/J* wild-type mice and performed left eye enucleation, followed by 5 days of dark rearing, and then 4 hours of light exposure (Fig. 4A). Upon collecting the brains, their slices were immunostained with the antibodies for c-Fos and Lamin B1, a component of the inner nuclear membrane (Fig. 4B). c-Fos staining confirmed appropriate stimulation for the cells in the ipsilateral visual cortex and the absence of stimulation in the contralateral visual cortex, in both young and aged brains. In the young brain, the nuclear shape observed by Lamin B1 exhibited more infolded than spherical morphology in the ipsilateral brain compared to the contralateral brain (Fig. 4B, C). However, the nuclear shapes in aged mice demonstrated an overall infolded appearance regardless of ipsilateral or contralateral condition (Fig. 4B, C). Also, the infolded nuclear state in the contralateral brain of aged mice was evaluated by circularity, area ratio, and nearest distance from the center of gravity analyses (Fig. 4D–F), and indicated nuclear shapes presenting a resting infolded state independent of any stimulation in the aged brain. This result suggests that brain aging induces nuclear infolding resembling the prior observed nuclear shapes in PD and HD brains ^7–11^, leading to the assumption that functional decline of the brain may be associated with infolded shapes of neuronal nuclei.

**Figure 4.**
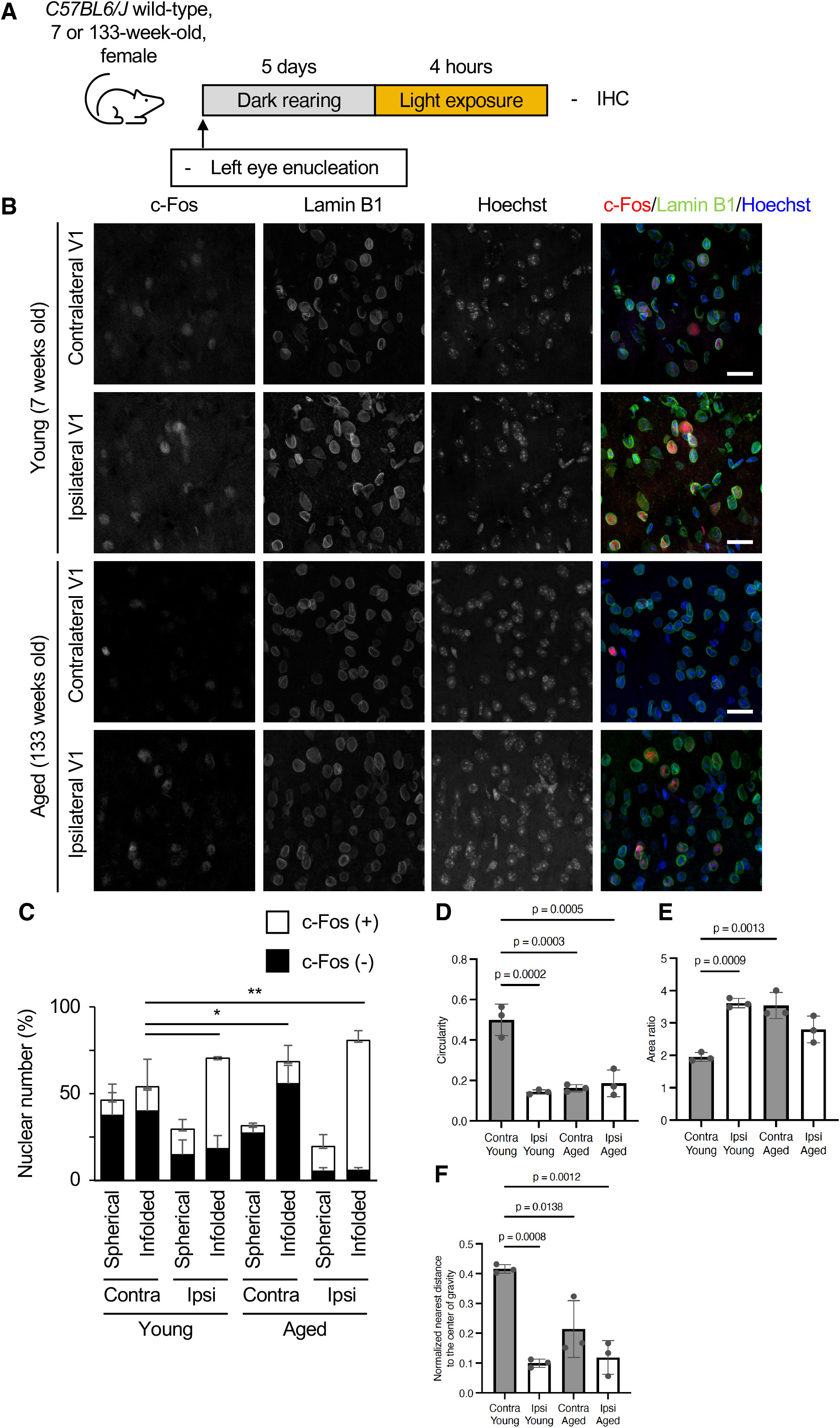
More infolded nuclei in the excitatory neurons in the aged primary visual cortex without visual stimulation. **(A)**7- or 133-week-old *C57BL6/J* wild-type mice were analyzed as in Fig. 1A. **(B)** Coronal sections of the brain were stained with the antibodies to c-Fos and Lamin B1. Nuclei were counterstained with Hoechst 33342. The images were obtained from layer 2/3 of the primary visual cortex. Scale bars, 20 μm. **(C)** Quantification of the proportion of c-Fos–positive and spherical or infolded nuclear shape in all layers of the primary visual cortex. 1,200 (young, contralateral), 966 (young, ipsilateral), 1,096 (aged, contralateral), and 1,176 (aged, ipsilateral) nuclei from three independent experiments were analyzed. (**D–F**) Quantification of circularity (D) or area ratio (E) value of Lamin B1 signals or nearest distance from the center of gravity (F) in the ipsilateral and contralateral visual cortex of young and aged mice. 202 (circularity, young, contralateral), 184 (circularity, young, ipsilateral), 143 (circularity, aged, contralateral), 147 (circularity, aged, ipsilateral), 179 (area ratio, young, contralateral), 173 (area ratio, young, ipsilateral), 126 (area ratio, aged, contralateral), 41 (area ratio, aged, ipsilateral), 189 (nearest distance, young, contralateral), 167 (nearest distance, young, ipsilateral), 139 (nearest distance, aged, contralateral), and 150 (nearest distance, aged, ipsilateral) nuclei from three independent experiments were analyzed. *p* values were determined by one-way ANOVA followed by Tukey’s multiple-comparison test.

### The nuclei in the aged brain have reduced nuclear shape dynamics compared to young nuclei

To reveal the temporal dynamics of aged nuclei, we performed time-lapse imaging of nuclear shape in upper layer neurons using *Nex-Cre;SUN1-GFP* mice that were more than 2 years old, as in Fig. 2 (Fig. 5A). Similar to the immunohistochemistry data (Fig. 4B), we found that more nuclei presented an infolded shape and the area ratio value was higher in aged mice compared to young mice even before stimulation (Fig. 5B, C, Supplementary Videos 5, 6). Additionally, the nuclear shape did not alter during stimulation and maintained a stable area ratio value throughout the imaging (Fig. 5B, D). The lack of nuclear shape alteration in excitatory neurons toward visual stimulation in aged mice suggests that the dynamics of nuclei morphology reduce with age.

**Figure 5.**
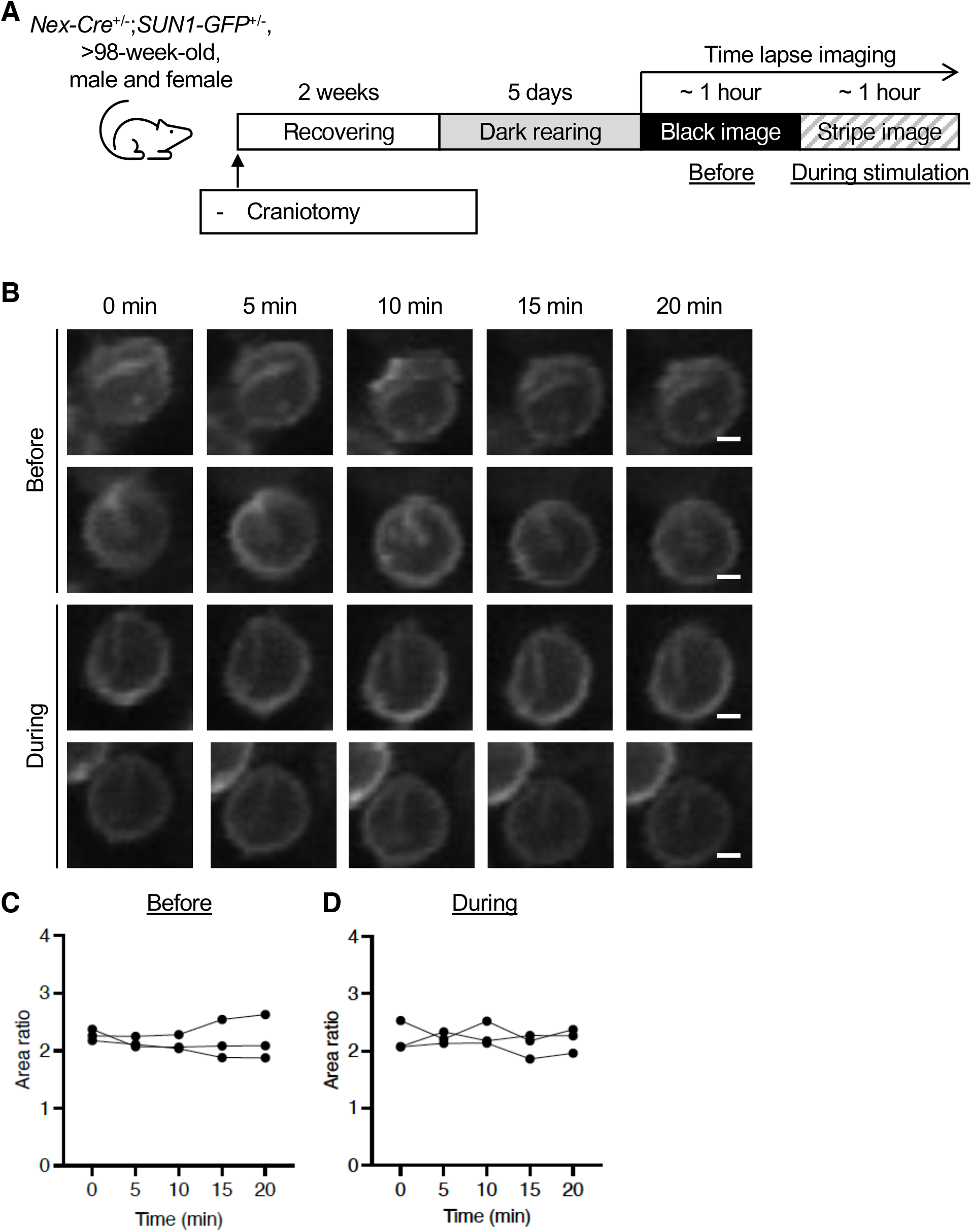
Less dynamic behavior of the nuclear shape in the aged brain. **(A)** *Nex-Cre;SUN1-GFP* mice older than 98 weeks old were analyzed as in Fig. 2A. **(B)** Representative images of GFP signals in “before” (upper two panels) and “during” (lower two panels) conditions. Scale bars, 2 μm. **(C, D)** Quantification of area ratio values at indicated time in “before” (C) or “during” (D) recording. 39 nuclei (C) and 33 nuclei (D) from three independent experiments were analyzed.

We next asked, why did the aged neurons show less frequent nuclear infolding events upon visual stimulation than young ones? Previous studies have shown that the nuclei of fibroblasts from patients with Hutchinson–Gilford progeria syndrome (HGPS), a premature aging disorder, were infolded and stiffer than those of the control cell line ^23–26^, suggesting an association between nuclear infolding and stiffness. In addition, nuclear stiffness changes during development and under conditions of cellular stress ^27,28^. Therefore, we assumed that stiffened nuclei would exhibit less frequent infolding upon aging when visually stimulated, and measured the stiffness of the isolated nuclei by AFM ^29^. Nuclei were isolated from the visual cortex of 12-to 16-week-old or 104-to 114-week-old *Nex-Cre;SUN1-GFP* mice using the Percoll density gradient centrifugation method ^30^, and only GFP-positive nuclei were subjected to AFM analysis (Fig. 6A). As a result, slope values in indentation force versus depth curve, representing stiffness, for aged nuclei were significantly larger than those of young nuclei (Fig. 6B, C). This result revealed for the first time that aged nuclei were stiffer than young nuclei during the natural aging process. Therefore, the stiffening of the nuclei may cause resistance to deformation and the less frequent infolding events observed in aged neurons upon visual stimulation.

**Figure 6.**
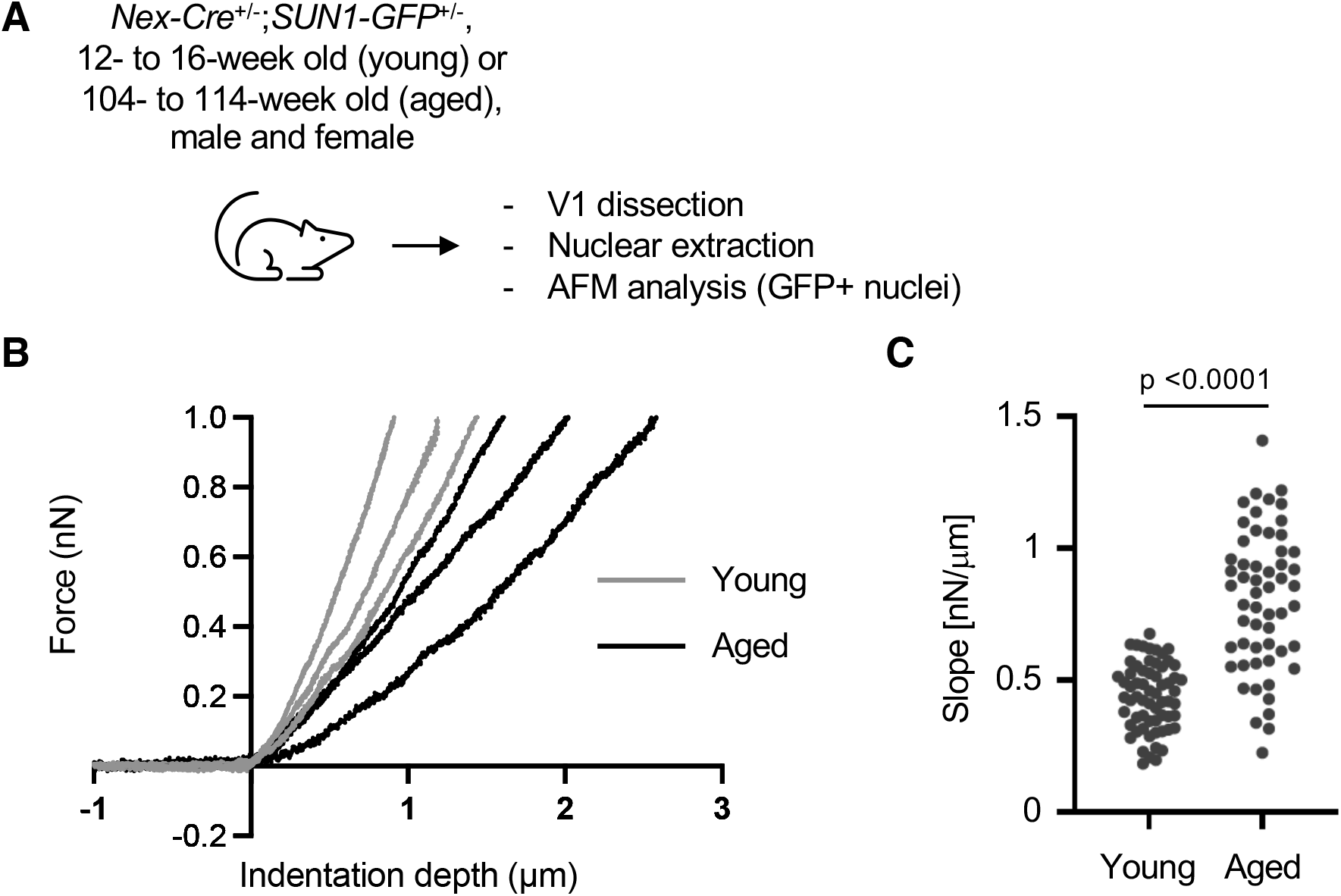
Stiffer nuclei of the excitatory neurons in the aged visual cortex. **(A)** Nuclei were isolated from the visual cortex of 12- to 16-week-old (young) or 104- to 114-week-old (aged) *Nex-Cre;SUN1-GFP* mice, and GFP-positive nuclei were analyzed by AFM. **(B)** Representative curves between indentation force and depth. Three curves each for young and aged nuclei are shown. **(C)** Beeswarm plots for the slope values in indentation force versus depth curve for nuclei. 63 young nuclei and 58 aged nuclei from three independent experiments were analyzed. *p* values were determined by two-tailed Welch’s t-test.

## Discussion

Nuclear infolding in neurons upon neuronal stimulation or neurodegenerative diseases has been reported in previous papers ^7–11,15,16^, but its regulatory mechanism and significance have not been fully understood. In this study, we found that an in vivo physiological stimulation through light stimulation of the visual cortex induced nuclear infolding in excitatory neurons, similar to exposure to a novel environment for the hippocampal neurons ^16^. Furthermore, time-lapse imaging of the nuclear shape of excitatory neurons in the upper layer showed the dynamic behavior of the nucleus during light stimulation. As indicated in a previous study, immunocytochemistry showed that inhibition of neuronal activity with tetrodotoxin (TTX) in a hippocampal neuronal culture decreases the infolded nuclei ^15^. Our time-lapse imaging of inhibition induced by light stimulation or MK-801 injection also revealed that nuclear infolding upon stimulation is a reversible process.

Treatment with MK-801 reduced the dynamic behavior, similar to the previous study using hippocampal slice culture ^15^. This suggests that calcium influx mediated by the NMDA receptor is upstream of nuclear infolding. In our imaging analysis, it took approximately 15 minutes of light stimulation for the dynamic behavior of the nucleus to begin. A previous study using hippocampal neuronal culture has shown the involvement of MAP kinase signaling in this process as MEK inhibitors, such as PD98059 and U0126, inhibited neuronal activity–dependent nuclear infolding ^15^. Upon KCl stimulation, MAP kinase, such as Erk, becomes phosphorylated and activated within 5 minutes ^22^. This is consistent with the idea that dynamic behavior and infolding of a neuronal nucleus are mediated by MAP kinase.

External stimuli induce the transcription of several genes in neurons and play an important role in the functional changes of neurons, such as neurite and synapse formation ^13,14^. Among these induced genes, a gene set called IEGs or rapid primary responsive genes (PRGs), including *c-Fos* and *Arc,* increased their expression levels within 20 minutes of visual stimulation and decreased after turning off the stimulation ^22^. This time course and reversibility are similar to the induction of nuclear infolding observed in the time-lapse imaging of this study. Considering that the components of the inner nuclear membrane, such as Lamin, are involved in gene transcription through regulating chromatin structure, the changes in the nuclear shape may contribute to the regulation of gene transcription. This is further supported by the previous work studying the role of the interaction between Satb2, a neuronal transcription factor, and Lemd2, a component of the inner nuclear membrane, in nuclear infolding upon neuronal activity ^16^. How then does nuclear infolding contribute to gene induction? One possibility is that nuclear infolding may allow rapid interactions between IEG loci and their regulatory elements. A previous Hi-C experiment using hippocampal neurons after kainic acid injection has shown the chromatin interactions at the gene loci of responsive genes, such as *c-Fos* and *BDNF,* within 1 hour ^31^. By contrast, the interaction between the nuclear membrane and the responsive gene loci, such as *Egr3* and *BDNF,* visualized by fluorescence in situ hybridization (FISH), has not been altered after bicuculline treatment in hippocampal neuronal culture ^32^. Therefore, the nuclear membrane may be required to be infolded in order for the regulatory element to interact with the gene loci while still bound to the nuclear membrane.

In the aged visual cortex of mice older than 2 years, the nuclei of excitatory neurons generally demonstrated a higher portion of infolded nuclei than the young visual cortex in non-stimulated conditions. In the previous study, the invagination of the nuclei of spiny projection neurons in the striatum of wild-type mice did not increase in aged mice (2 years old) compared to young mice (2 months old) ^9^. This discrepancy may be caused by the difference in the neuronal type, brain area, and harvesting condition of mice. What is the cause for the higher population of infolded nuclei in aged excitatory neurons? Long-term stimulation (40 hours) of hippocampal neurons by bicuculline induces a more stable infolded nuclear shape after TTX treatment compared to short-term stimulation (1 hour) ^15^. This suggests that more stimulation induces the fixed nuclear-infolded shape. Aged excitatory neurons have naturally undergone repeated neuronal activity exposure and this may cause the irreversible infolded nuclear shape and the dysfunction of neuronal nuclei. In addition, the stiffer nuclei of aged neurons, observed in this study, can contribute to the irreversible nuclear shape. Consistently, our time-lapse imaging of nuclei of aged excitatory neurons revealed a decrease in dynamic behavior compared to those of young ones. We currently do not know the relationship between repetitive neuronal activity and nuclear shape dynamics, such as stiffness and frequency of morphological change; however, their additional or synergistic contribution may induce nuclear infolding in the aged brain.

The more infolded nuclear shape in the aged visual cortex in this study is reminiscent of the nuclear shape in PD and HD ^7,8,10,11^. Since both PD and HD are age-associated neurodegenerative disorders, our results indicate that an increase in infolded nuclei is associated with the decline of neuronal functions. Previous studies have shown defects in the compartmentalization of infolded nuclei (Shani et al. 2019; Alcalá-Vida et al. 2021; Gasset-Rosa et al. 2017) and suggest that leakiness or abnormal nucleocytoplasmic transport of mRNA and proteins cause a functional decline of aged neurons. Indeed, previous studies revealed leaky nuclear membranes of aged brain cells and defects of protein import into the nucleus in aged human fibroblast ^33,34^. In another study, micropipette aspiration of the nuclear membrane of embryonic stem cells (ESCs), differentiated neural progenitor cells, and tissue fibroblast revealed a more deformable nuclear membrane of ESCs compared to differentiated cells ^27^. ESCs have more plasticity for lineage choice and gene transcription (Meshorer and Misteli, 2006). Therefore, it is supposed that deformable nuclear membrane contributes to plasticity. Younger neurons also have more plasticity and are more adaptable to functional changes ^35,36^. Therefore, the less dynamic and stiffer nucleus in aged neurons may be related to their decrease in plasticity through regulating the events induced by neuronal activity, such as induction of IEGs.

## Material and Methods

### Animals

*ICR* or *C57BL/6J* mice (CLEA Japan) were used for the study of wild-type mice. *Nex-Cre* mice were provided by Klaus-Armin Nave (Max Planck Institute), and *SUN1-GFP* mice (#021039) were obtained from The Jackson Laboratory. The age of the mice is indicated in the figure legends. All mice were maintained in a temperature- and relative humidity–controlled (23 ± 3°C and 50 ± 15%, respectively) environment with a normal 12-hour light and 12-hour dark cycle. Two to six mice were housed per sterile cage (Innocage, Innovive; or Micro BARRIER Systems, Edstrom Japan) with chips (PALSOFT, Oriental Yeast; or PaperClean, SLC Japan), and irradiated food (CE-2, CLEA Japan) and filtered water available ad libitum. All animals were maintained and studied according to protocols approved by the Animal Care and Use Committee of The University of Tokyo.

### Enucleation and dark rearing in mice for neuronal activity manipulation

The eye removal was performed under 2%isoflurane. The mouse was placed on a little pedestal. A tweezer clipped the optical nerve behind the eye, and then the eye was removed. The wound was glued via tissue adhesive application from Vetbond. After the left eye removal and the mouse’s recovery, they were exposed to 5 days of darkness within a box covered by a dark sheet to obtain the neuronal activity of the visual cortex to its lowest baseline.

### RT-qPCR analysis

The visual cortex was dissected with a stainless steel 0.5 mm sagittal brain matrix for rodents. The brain was first cut at 1.5 mm and 3.5 mm from the midline. Then, the visual cortex was located over one-third of the posterior part of the hippocampus. The dissected visual cortex was put into RNAiso plus (Takara-Bio), and total RNA was precipitated. The amounts of obtained RNA were measured via NanoDrop One (Thermo Fisher Scientific), and 0.5 μg of the RNA was subjected to reverse transcription with the use of ReverTra Ace qPCR RT Master Mix with gDNA Remover (Toyobo). The resultant cDNA was subjected to real-time PCR analysis in a LightCycler 480 or LC96 instrument (Roche) with Thunderbird SYBR qPCR mix (Toyobo) or QuantiNova SYBR green PCR kit (Qiagen). The amount of each target mRNA was normalized by that of *GAPDH* mRNA. Primer sequences are provided in Supplementary Table 1.

### Immunohistochemistry

Immunohistochemistry was performed as previously described ^37^. Mice were injected intraperitoneally with pentobarbital solution (Nakarai Tesque) for anesthesia. Cold PBS was flushed via a 30G needle, followed by clod 4% paraformaldehyde (PFA) in phosphate-buffered saline (PBS) on the left heart ventricle. After perfusion, the brains were removed and soaked overnight in 4% PFA on a rotator, and then they were incubated with 10%, 20%, and 30% sucrose in PBS and embedded with O.C.T. compound (Sakura Finetek Japan) at −80°C. Sixteen-micrometer coronal sections of embedded brains were prepared by a cryostat. For staining, samples were exposed to antigen retrieval (10 minutes at 105°C in 1 x TRS (DAKO)). Then the samples were exposed to Tris-buffered saline containing 0.1% Triton X-100 and 3% bovine serum albumin (blocking buffer) for 2 hours at room temperature, incubated first overnight at 4°C with primary antibodies (Supplementary Table 2) in blocking buffer and then for 2 hours at room temperature with Alexa Fluor–conjugated secondary antibodies (1: 1000, Thermo Fisher Scientific) in blocking buffer, and mounted in Mowiol (Calbiochem). All images were taken with Leica SP5 or Zeiss LSM 880 as a z-stack image to include the entire nucleus. For analysis of the images, ImageJ software (National Institute of Health) was used. Nuclei for which the entire image was taken were randomly selected from maximum projection images and analyzed for subjective counting of infolded or spherical nuclei, circularity, and nearest distance from the center of gravity. For subjective counting, infolded nuclei were defined by shaped or patterned signals inside the nucleus or by the outer shape being strongly invaginated. For circularity, area ratio, and nearest distance from the center of gravity, the GFP or Lamin B1 signals were binarized (Fig. 2D). The inner nuclear shape (gray in Fig. 2D) was used for the calculation of circularity. After determining the center of gravity on the basis of the outer nuclear shape, the distance from this center to the closest signal of GFP or Lamin B1 was normalized with the apparent radius.

### Craniotomy and viral injection into the visual cortex

*Nex-Cre;SUN1-GFP* mice were prepared for in vivo imaging as described previously ^38,39^. Under stereotaxis conditions, which were performed under constant 2% isoflurane exposure, the skin over the skull was removed, and any muscle in proximity was cleanly cut off to ensure a dry area on which the plate could be positioned and glued by the Super-bond dental cement (Sun Medical). A custom-made metal head plate was glued on the skull, and the area of interest was located at the caudal region around 4 mm from Lambda in the partial bone, the visual cortex. The craniotomy was made over the entire visual cortex, and the dura (thin protective layer on the brain) was carefully removed. Five-hundred nanoliters of the virus, AAV1-Syn-Flex-NES-jRGECOSa-WPRE-SV40 (Plasmid #100854, Addgene) ^40^, in saline was directly injected into the cortex at a speed of 1 nL/s by Nanoject III (Drummond), while covered by artificial CSF. Afterward, the window was closed with glass slips (5.5 mm and 4 mm glued onto each other) and use of clear glue and dental cement (ADFA, Shofu). The closed window was further covered for protection with dental silicone (Shofu). After approximately 2 weeks, the window was checked for its clarity and covered again with dental silicone. Five days prior to the recordings, the mice were put into a dark rearing box.

### Microinjection of inhibitor

The skin was opened and removed to insert a pinhole with a 30G needle onto the visual cortex (the caudal region around 4 mm from Lambda in the partial bone) in the mice under 2% isoflurane anesthesia in stereotaxis fixation. Five-hundred nanoliters of 100 mM MK-801 in saline was injected at a speed of 1 nL/s into the ipsilateral hemisphere by the Nanoject III. The right visual cortex, as a control, received a saline injection. The skin was stitched closed after injection.

### Two-photon microscopy of the visual cortex of *Nex-Cre;SUN1-GFP* mice

Approximately 2 weeks after the craniotomy, the mice were exposed to darkness for 5 days in a dark rearing box and seated on a heat block during the entire time of recording. During anesthetized recordings, mice were sedated with chlorprothixene (0.3–0.8 mg/kg; Sigma-Aldrich) and isoflurane (0.8%). The recordings were done with NIS-Elements AR (Nikon) using a two-photon microscope (Nikon A1MP) equipped with a mode-locked Ti:sapphire laser (Mai Tai Deep See; Spectra Physics). The excitation light was focused with a 25x objective (Nikon Plan Apo, NA 1.10). GFP and jRGECO signals were excited at 960 nm, and the emission was filtered at 517–567 nm and 570–625 nm, respectively. A square region of the cortex (512 x 512 pixels, approximately 330 μm on a side) was imaged at 2 frames/min. Images were obtained from layers 2/3 (200–300 μm from the pia). Visual stimuli were generated using custom-written programs in PsychoPy (Peirce, 2007). The stimulus presentation was synchronized with the frame acquisition of images using a counter board (NI USB-6501; National Instruments). We positioned a 32-inch LCD monitor 18 cm from the right eye of each mouse. Drifting rectangle strips of different angles were presented as waves in a randomized manner. For the conversion of the recordings, Matlab R2020b (version 4 9.9) was used. Note that we did not use the jRGECO signals for further analysis because of insufficient image quality. For analysis of the images, ImageJ software was used. Nuclei whose centers were taken in the z-axis direction were randomly selected for analysis of area ratio.

### Nuclear isolation

Nuclear isolation was performed as previously described ^30^. The visual cortex isolated as in RT-qPCR analysis was homogenized using a syringe with 23G and 27G needles, sequentially, in 500 μL of 54% Percoll in homogenizing buffer (50 mM Tris-HCl pH 7.4, 25 mM KCl, 5 mM MgCl2, and 250 mM sucrose) on ice. The solution was mixed with 10% NP-40 (final concentration, 0.1%), left on ice for 15 minutes, and mixed with 500 μL of homogenizing buffer. Then, Percoll gradient was prepared in the following order: 100 μL of 35% Percoll in homogenizing buffer on the bottom layer, 200 μL 31% Percoll in homogenizing buffer, and 1 mL of homogenate (27% Percoll) on the top layer. The tube was centrifuged at 20,000 x g for 10 minutes at 4°C. After removing the debris on the top layer, nuclei on the bottom layer were transferred into another tube.

### Mechanical testing for isolated nuclei using AFM

Cell nuclei of excitatory neurons in the visual cortex of *Nex-Cre;SUN1-GFP* mice were attached to the Poly-D-Lysine (PDL)-coated Petri dishes (World Precision Instrument, FD5040-100) and subjected to AFM-based mechanical testing ^41^. In this study, we utilized JPK BioAFM NanoWizard 3 (Bruker Nano GmbH), which was mounted on an epi-fluorescence microscope (IX81; Evident). The spring constants of AFM cantilevers (qp-BioAC CB2; Nanoworld AG) were calibrated based on the thermal noise method ^42^. Cell nuclei of excitatory neurons were fluorescently identified with GFP. AFM mechanical testing for the GFP-positive nuclei was conducted with the following settings; the piezo displacement speed of 6 μm/s and the sampling rate of 4,000 Hz. Based on the indentation force (*F*) versus depth curve obtained by mechanical testing, we estimated slope [nN/μm] by linear regression for the sample points within a force range (500 pN ≤ *F* ≤ 1,000 pN), as previously described ^43,44^.

### Statistical analysis

Data are presented as means ± s.d. and were compared with two-tailed Welch’s *t*-test or by analysis of variance (ANOVA) followed by Tukey’s multiple-comparison test. A p value of < 0.05 was considered statistically significant.

## Acknowledgments

We thank Klaus-Armin Nave (Max Planck Institute) for providing *Nex-Cre* mice, Jeremy Nathans (John Hopkins Medical School) for providing *SUN1-GFP* mice, Miki Bundo (Kumamoto University) for the method of nuclear isolation, Eetsuko Ogawara for technical assistance, Ayuko Hoshino (Tokyo Institute of Technology) for critical review of this manuscript, and members of the Gotoh and Kishi laboratories for their discussions. This research was supported by MEXT Scholarship for T.F. AMED-CREST (22gm1310004 to Y.G.), AMED-PRIME (JP22gm6110021 to Y.K.), and MEXT/JSPS KAKENHI (22H00431 to Y.G.; 20H03179, 21H00242, and 22K19823 to Y.K.), by the Takeda Science Foundation, by the Mochida Memorial Foundation for Medical and Pharmaceutical Research, and by the SECOM Science and Technology Foundation.

## Author contributions

T.F. and Y.K. designed the study and wrote the manuscript. T.F., T.M., K.M., T.K., N.N., A.S., and Y.K. performed the experiments and analyzed the data. T.M. and K.O. assisted with time-lapse imaging. T.A. and M.K. assisted with AFM analysis. Y.G. and Y.K. supervised the study. All authors contributed to the revision and approved the final version of the manuscript.

## Figure legends

**Supplementary figure 1.**
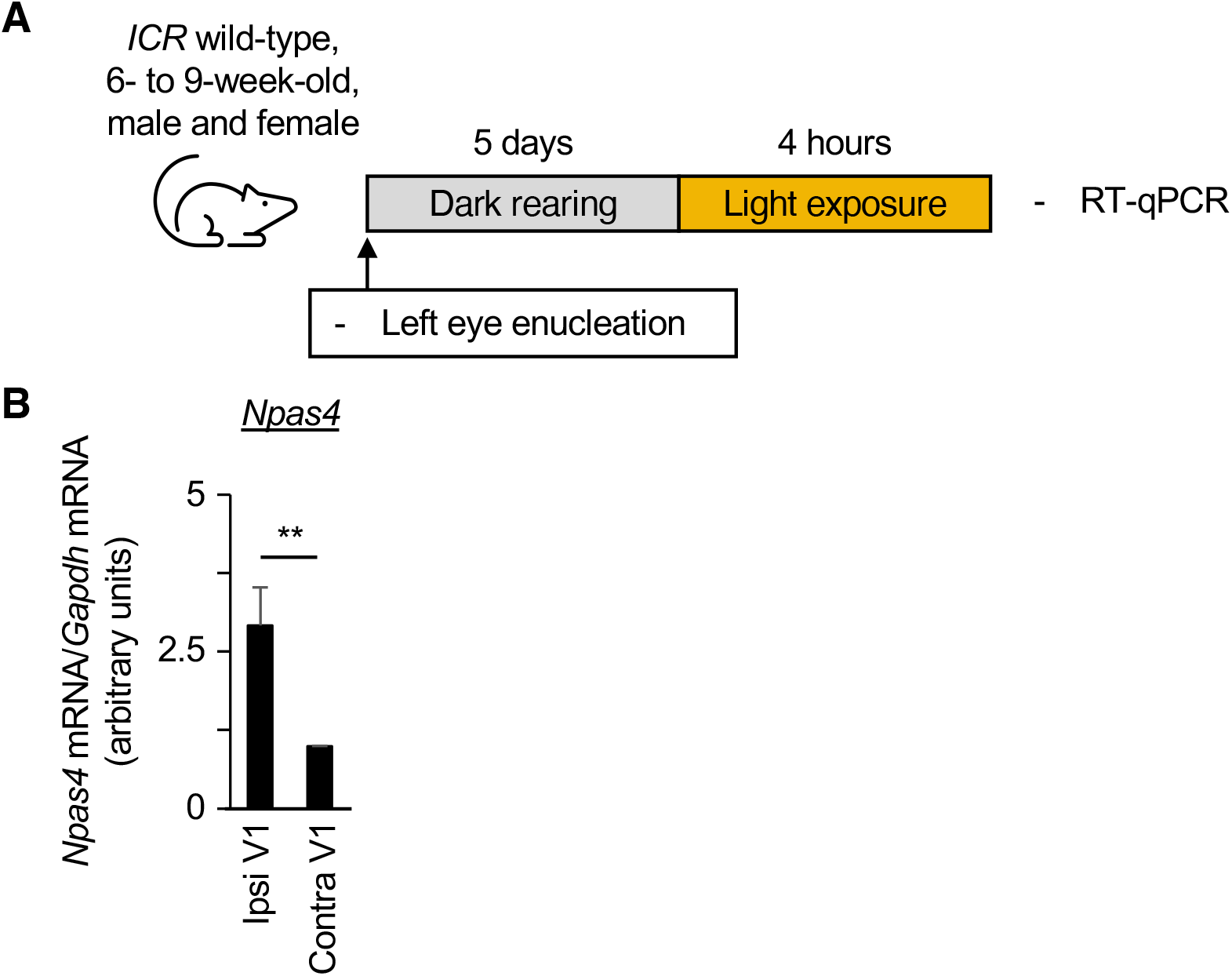
Expression changes of an IEG upon visual stimulation after enucleation, dark rearing, and light exposure. **(A)** Left eyes of 6- to 9-week-old *ICR* wild-type mice were enucleated, and the mice were kept in a dark room for 5 days. They were stimulated by room light for 4 hours and subjected to reverse transcription and quantitative polymerase chain reaction (RT-qPCR). **(B)** RT-qPCR analysis of relative *Npas4* mRNA abundance (normalized by the amount of *Gapdh* mRNA) in the primary visual cortex was performed. Data are means ± s.d., averaged values for four mice. \*\**p* < 0.01 (two-tailed Wlech’s t-test).

**Supplementary figure 2.**
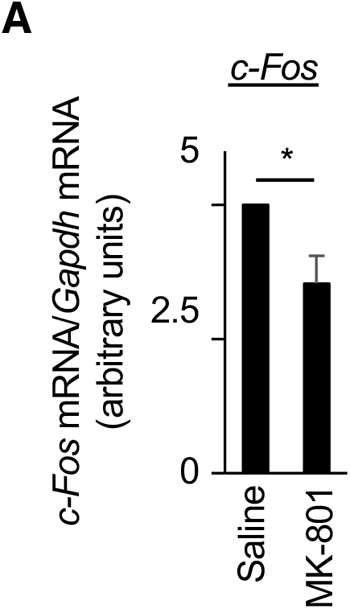
The changes in the expression level of an IEG after MK-801 injection. **(A)** RT-qPCR analysis of relative *c-Fos* mRNA abundance (normalized by the amount of *Gapdh* mRNA) in the primary visual cortex after injection of saline or MK-801 was performed. Data are means ± s.d., averaged values for four mice. \**p* < 0.05 (two-tailed Welch’s t-test).

**Supplementary Video 1**

The nuclear shape of excitatory neurons in the visual cortex of 6- to 9-week-old *Nex-Cre;SUN1-GFP* mouse before stimulation. This video is played at 120× speed.

**Supplementary Video 2**

The nuclear shape of excitatory neurons in the visual cortex of 6- to 9-week-old *Nex-Cre;SUN1-GFP* mouse during stimulation. The nuclei shown in Figure 2 are indicated as yellow arrows. This video is played at 120× speed.

**Supplementary Video 3**

The nuclear shape of excitatory neurons in the visual cortex of 6- to 9-week-old *Nex-Cre;SUN1-GFP* mouse after stimulation. This video is played at 120× speed.

**Supplementary Video 4**

The nuclear shape of excitatory neurons in the visual cortex of 6- to 9-week-old *Nex-Cre;SUN1-GFP* mouse after MK-801 injection. This video is played at 120× speed.

**Supplementary Video 5**

The nuclear shape of excitatory neurons in the visual cortex of *Nex-Cre;SUN1-GFP* mice older than 98 weeks before stimulation. This video is played at 120× speed.

**Supplementary Video 6**

The nuclear shape of excitatory neurons in the visual cortex of *Nex-Cre;SUN1-GFP* mice older than 98 weeks during stimulation. This video is played at 120× speed.

**Supplementary Table 1.**
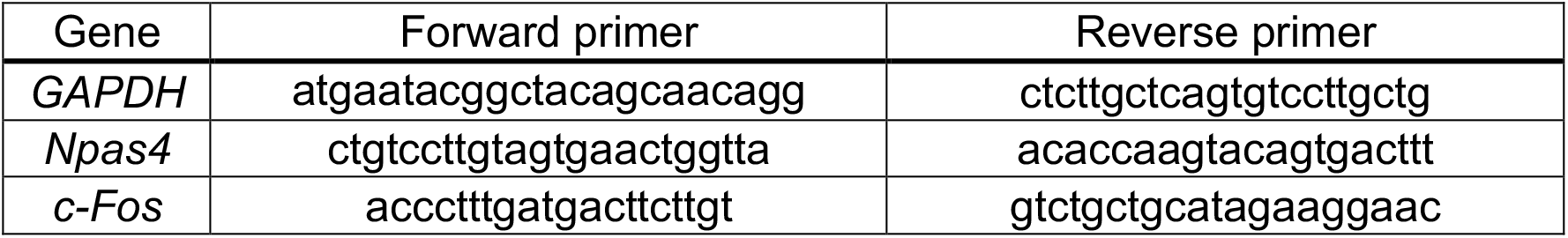
Primer sequences for RT-qPCR.

**Supplementary Table 2.** Antibodies for immunohistochemistry.

| Antigen | Manufacturer | Serial Number | Concentration |
| --- | --- | --- | --- |
| GFP | Abcam | ab13970 | 1:2000 |
| c-Fos | Santa Cruz | sc-271243 | 1:100 |
| Lamin B1 | Abcam | ab16048 | 1:250 |

## References

1. Camandola, S. & Mattson, M. P. Brain metabolism in health, aging, and neurodegeneration. Embo J 36, 1474–1492 (2017).

2. Hof, P. R. & Morrison, J. H. The aging brain: morphomolecular senescence of cortical circuits. Trends Neurosci 27, 607–613 (2004).

3. Hou, Y. et al. Ageing as a risk factor for neurodegenerative disease. Nat Rev Neurol 15, 565–581 (2019).

4. Scaffidi, P. & Misteli, T. Lamin A-dependent nuclear defects in human aging. Science 312, 1059–1063 (2006).

5. Heckenbach, I. et al. Nuclear morphology is a deep learning biomarker of cellular senescence. Nat Aging 2, 742–755 (2022).

6. Ragnauth, C. D. et al. Prelamin A Acts to Accelerate Smooth Muscle Cell Senescence and Is a Novel Biomarker of Human Vascular Aging. Circulation 121, 2200–2210 (2010).

7. Shani, V. et al. Physiological and pathological roles of LRRK2 in the nuclear envelope integrity. Hum Mol Genet 28, 3982–3996 (2019).

8. Liu, G.-H. et al. Progressive degeneration of human neural stem cells caused by pathogenic LRRK2. Nature 491, 603–607 (2012).

9. Chen, X. et al. Parkinson’s disease-related Leucine-rich repeat kinase 2 modulates nuclear morphology and genomic stability in striatal projection neurons during aging. Mol Neurodegener 15, 12 (2020).

10. Alcalá-Vida, R. et al. Neuron type-specific increase in lamin B1 contributes to nuclear dysfunction in Huntington’s disease. Embo Mol Med 13, e12105 (2021).

11. Gasset-Rosa, F. et al. Polyglutamine-Expanded Huntingtin Exacerbates Age-Related Disruption of Nuclear Integrity and Nucleocytoplasmic Transport. Neuron 94, 48–57.e4 (2017).

12. Wong, R. O. L. & Ghosh, A. Activity-dependent regulation of dendritic growth and patterning. Nature Reviews Neuroscience 3, 803–812 (2002).

13. Greer, P. L. & Greenberg, M. E. From synapse to nucleus: calcium-dependent gene transcription in the control of synapse development and function. Neuron 59, 846–860 (2008).

14. Flavell, S. W. & Greenberg, M. E. Signaling Mechanisms Linking Neuronal Activity to Gene Expression and Plasticity of the Nervous System. Annu Rev Neurosci 31, 563–590 (2008).

15. Wittmann, M. et al. Synaptic activity induces dramatic changes in the geometry of the cell nucleus: interplay between nuclear structure, histone H3 phosphorylation, and nuclear calcium signaling. Journal of Neuroscience 29, 14687–14700 (2009).

16. Feurle, P. et al. SATB2-LEMD2 interaction links nuclear shape plasticity to regulation of cognition-related genes. The EMBO Journal e103701 (2020) doi:10.15252/embj.2019103701.

17. Mo, A. et al. Epigenomic Signatures of Neuronal Diversity in the Mammalian Brain. Neuron 86, 1369–1384 (2015).

18. Goebbels, S. et al. Genetic targeting of principal neurons in neocortex and hippocampus of NEX-Cre mice. Genesis 44, 611–621 (2006).

19. Lyckman, A. W. et al. Gene expression patterns in visual cortex during the critical period: synaptic stabilization and reversal by visual deprivation. Proceedings of the National Academy of Sciences 105, 9409–9414 (2008).

20. Hensch, T. K. Critical period plasticity in local cortical circuits. Nature Reviews Neuroscience 6, 877–888 (2005).

21. Hooks, B. M. & Chen, C. Critical Periods in the Visual System: Changing Views for a Model of Experience-Dependent Plasticity. Neuron 56, 312–326 (2007).

22. Tyssowski, K. M. et al. Different Neuronal Activity Patterns Induce Different Gene Expression Programs. Neuron 98, 530–546.e11 (2018).

23. Verstraeten, V. L. R. M., Ji, J. Y., Cummings, K. S., Lee, R. T. & Lammerding, J. Increased mechanosensitivity and nuclear stiffness in Hutchinson–Gilford progeria cells: effects of farnesyltransferase inhibitors. Aging Cell 7, 383–393 (2008).

24. Csoka, A. B. et al. Novel lamin A/C gene (LMNA) mutations in atypical progeroid syndromes. J Med Genet 41, 304 (2004).

25. Eriksson, M. et al. Recurrent de novo point mutations in lamin A cause Hutchinson–Gilford progeria syndrome. Nature 423, 293–298 (2003).

26. Goldman, R. D. et al. Accumulation of mutant lamin A causes progressive changes in nuclear architecture in Hutchinson–Gilford progeria syndrome. Proc National Acad Sci 101, 8963–8968 (2004).

27. Pajerowski, J. D., Dahl, K. N., Zhong, F. L., Sammak, P. J. & Discher, D. E. Physical plasticity of the nucleus in stem cell differentiation. Proceedings of the National Academy of Sciences of the United States of America 104, 15619–15624 (2007).

28. Nava, M. M. et al. Heterochromatin-Driven Nuclear Softening Protects the Genome against Mechanical Stress-Induced Damage. Cell 181, 800–817.e22 (2020).

29. Newberg, J. et al. Isolated nuclei stiffen in response to low intensity vibration. J Biomech 111, 110012 (2020).

30. Bundo, M., Kato, T. & Iwamoto, K. Epigenetic Methods in Neuroscience Research. Neuromethods 115–123 (2016) doi:10.1007/978-1-4939-2754-8_7.

31. Fernandez-Albert, J. et al. Immediate and deferred epigenomic signatures of in vivo neuronal activation in mouse hippocampus. Nat Neurosci 22, 1718–1730 (2019).

32. Noguchi, A. et al. Decreased Lamin B1 Levels Affect Gene Positioning and Expression in Postmitotic Neurons. Neurosci Res 173, 22–33 (2021).

33. D’Angelo, M. A., Raices, M., Panowski, S. H. & Hetzer, M. W. Age-dependent deterioration of nuclear pore complexes causes a loss of nuclear integrity in postmitotic cells. Cell 136, 284–295 (2009).

34. Pujol, G., Söderqvist, H. & Radu, A. Age-associated reduction of nuclear protein import in human fibroblasts. Biochem Bioph Res Co 294, 354–358 (2002).

35. Katz, L. C. & Crowley, J. C. Development of cortical circuits: Lessons from ocular dominance columns. Nat Rev Neurosci 3, 34–42 (2002).

36. Berardi, N., Pizzorusso, T. & Maffei, L. Critical periods during sensory development. Curr Opin Neurobiol 10, 138–145 (2000).

37. Eto, H. et al. The Polycomb group protein Ring1 regulates dorsoventral patterning of the mouse telencephalon. Nature Communications 11, 5709 (2020).

38. Murakami, T., Yoshida, T., Matsui, T. & Ohki, K. Wide-field Ca2+ imaging reveals visually evoked activity in the retrosplenial area. Front Mol Neurosci 08, 20 (2015).

39. Ohki, K. & Reid, R. C. In Vivo Two-Photon Calcium Imaging in the Visual System. Cold Spring Harb Protoc 2014, pdb.prot081455 (2014).

40. Dana, H. et al. Sensitive red protein calcium indicators for imaging neural activity. Elife 5, e12727 (2016).

41. Maki, K. et al. Mechano-adaptive sensory mechanism of α-catenin under tension. Sci Rep-uk 6, 24878 (2016).

42. Butt, H.-J. & Jaschke, M. Calculation of thermal noise in atomic force microscopy. Nanotechnology 6, 1 (1995).

43. Ichijo, R. et al. Vasculature atrophy causes a stiffened microenvironment that augments epidermal stem cell differentiation in aged skin. Nat Aging 2, 592–600 (2022).

44. Maki, K., Han, S.-W. & Adachi, T. β-Catenin as a Tension Transmitter Revealed by AFM Nanomechanical Testing. Cell Mol Bioeng 8, 14–21 (2015).

